# Modular and integrative activity reporters enhance biochemical studies in the yeast ER

**DOI:** 10.1101/2023.07.12.548713

**Authors:** Samantha G Martinusen, Ethan W Slaton, Sage E Nelson, Marian A Pulgar, Julia T Besu, Cassidy F Simas, Carl A Denard

## Abstract

The yeast endoplasmic reticulum sequestration and screening (YESS) system is a generalizable platform that has become highly useful to investigate post-translational modification enzymes (PTM-enzymes). This system enables researchers to profile and engineer the activity and substrate specificity of PTM-enzymes and to discover inhibitor-resistant enzyme mutants. In this study, we expand the capabilities of YESS by transferring its functional components to integrative plasmids. The YESS integrative system yields uniform protein expression and protease activities in various configurations, allows one to integrate activity reporters at two independent loci and to split the system between integrative and centromeric plasmids. We characterize these integrative reporters with two viral proteases, Tobacco etch virus (TEVp) and 3-chymotrypsin like protease (3CL^pro^), in terms of coefficient of variance, signal-to-noise ratio and fold-activation. Overall, we provide a framework for chromosomal-based studies that is modular, enabling rigorous high-throughput assays of PTM-enzymes in yeast.

## Introduction

Enzymes that catalyze post-translational modifications (PTM-enzymes), including proteases (López-Otín and Bond 2008), kinases (Cheng, Qi et al. 2011), methyltransferases (Petrossian and Clarke 2011) and acetyltransferases (Kori, Sidoli et al. 2017) among others, play central roles in regulating the functional proteome. Aside from their biological roles, these enzymes, particularly proteases, protein ligases and several amino acid side-chain modification enzymes, can be leveraged for transformative biotechnological and biomedical applications (Barber and Rinehart 2018). Thus, a critical demand in this space is to develop genetically tractable and scalable high-throughput screening and selection platforms to engineer, profile and interrogate the substrate specificity and activity of PTM-enzymes. Among available technologies (Sellamuthu, Shin et al. 2011, Guerrero, O’Malley et al. 2016, Ravikumar, Arzumanyan et al. 2018, Meister, Hendrikse et al. 2019, Sanchez and Ting 2020, Blum, Liu et al. 2021), the yeast endoplasmic reticulum (ER) sequestration and screening (YESS) system has rapidly become an important platform to study and engineer PTM-enzymes. Mainly, YESS is compatible with proteases, tyrosine kinases and methyltransferases, while most other platforms who are largely restricted to proteases (Guerrero, O’Malley et al. 2016, Pethe, Rubenstein et al. 2019, Li, Leier et al. 2020, Zhou, Li et al. 2020, Blum, Liu et al. 2021). Furthermore, YESS allows the user to measure enzyme activities directly on the cell surface, rather than transcriptional signal amplification (Sanchez and Ting 2020). YESS is an all-in-one platform that allows one to engineer and profile the substrate specificity of PTM-enzymes while also enabling high-throughput drug screening deep mutational scanning assays (Yi, Gebhard et al. 2013, Yi, Taft et al. 2015, Li, Yi et al. 2017, Ezagui, Russell et al. 2022).

In a typical YESS plasmid, two gene cassettes, one harboring a PTM-enzyme and the other one or more of its substrates, are targeted to the yeast ER via an 5’-end ER signal peptide. The substrate cassette contains one or more DNA-encoded enzyme substrates flanked by epitope tags (i.e. FLAG and HA) fused to the 3’-end of aga2. Therefore, the substrate cassette will display on the yeast surface upon induction. The two gene cassettes can be driven by a bidirectional galactose promoter or can be tuned with orthogonal inducers. In addition, unique endoplasmic retention signals (ERS) placed at the 3’-end of each gene cassette dictate how strongly each polypeptide will interact with ER receptors. The strength of the ERS is dictated by its sequence (KDEL, FEHDEL, WEHDEL), thus enabling one to tune enzyme: substrate interaction time in the ER. As the substrate cassette migrates to the yeast membrane surface, enzyme-catalyzed modifications can be visualized using flow cytometry using fluorescently-labeled antibodies against flanking epitope tags or specific to the modified substrates. Since its inception, YESS has enabled several exciting advances in PTM-enzyme studies. YESS has been used to engineer TEVp with altered specificity and increased catalytic efficiency (Yi, Gebhard et al. 2013). It has been combined with Next-Gen generation (YESS-NGS) to profile the substrate specificity of host and viral proteases and discover drug-resistant kinase mutants (Li, Yi et al. 2017, Taft, Georgeon et al. 2019). Additionally, it has been used in conjunction with machine learning to predict the substrate preference and binding energy of hepatitis C virus (HCV) protease (Pethe, Rubenstein et al. 2019). More recently, YESS was used to assay histone acetyltransferase activities (Waldman, Rao et al. 2021) and crosstalk and to recapitulate mammalian phosphorylation cascades (Ezagui, Russell et al. 2022).

In theory, any PTM-enzyme whose activity can be detected on the yeast surface can be assayed via yeast ER sequestration, making YESS a generally applicable platform for studying PTM-enzymes in a heterologous host. Yet, despite these advances, the YESS system still requires significant optimizations to obtain desirable PTM-enzyme activities in the ER. One desirable feature is a high dynamic range. To achieve this dynamic range, one can manipulate enzyme: substrate ratios in the ER by modulating enzyme transcription levels. By finding the right combination of ER retention signals on the enzyme and substrate sides, one can modulate enzyme: substrate contact time, thus influencing enzyme kinetics post-translationally. On one hand, an enzyme with poor folding kinetics or an enzyme that exhibits low catalytic efficiency (∼10^2^ M^-1^s^-1^) on a chosen substrate, may require the strongest ERS on the substrate and enzyme cassettes (Mei, Zhai et al. 2017). On the other hand, removing the ERS from one of both cassettes selects for enzymes and substrates with the highest catalytic efficiencies. With this versatility, there is a need to achieve homogeneous protein expression, which translates into a high signal-to-noise ratio (SNR) of enzymatic activity in a cell population. On the yeast surface, homogeneous enzyme activity results in fluorescence signals with low statistical variance (Elowitz, Levine et al. 2002, Yi, Gebhard et al. 2013, Lee, DeLoache et al. 2015). Here, low statistical variance is particularly important in PTM-enzyme engineering and substrate profiling, allowing one to differentiate between clones with incremental differences in activities.

In a previous report, we modified YESS to quantify enzyme catalytic turnovers (YESS 2.0) more accurately (Denard, Paresi et al. 2021). Compared to the original system, YESS 2.0 allows one to control enzyme: substrate stoichiometries by titrating enzyme transcription with a hormone-inducible promoter. This iteration of YESS allows for engineering proteases with increased catalytic efficiency. Yet, because the two gene cassettes are plasmid-based, YESS generally suffers from high variance in protease activities, particularly at low enzyme activity settings. To address this, we have developed a suite of integrative YESS plasmids that achieve significantly low variability in protease activity compared to the current centromeric plasmid-based systems. We show that integrating protease activity reporters improves the visualization and quantification of proteolytic activity without impeding enzymatic activity. We further validate this system for use in directed evolution campaigns through a comparison of display levels when one cassette is integrated, and the other remains on a plasmid. For each configuration, we quantify the coefficient of variance (CV_HA_), fold activation and protease activity levels (D_dB,HA_) (Giesecke, Feher et al. 2017). In each of these applications, we show that the integration of activity reporters can improve the uniformity of gene transcription and D_dB,HA_ using two protease examples. We also provide a means to partially integrate YESS components to streamline library screening. Together our integrative YESS plasmids provide a robust platform to observe and quantify PTM-enzyme activity in yeast. We have shared these plasmids through Addgene with the research community, hoping to expand the use of ER sequestration across various PTM-enzyme assays.

## Results and Discussion

### Building integrative activity reporter plasmids

To begin, we built integrative enzyme activity reporters that mirror the plasmid-based systems published previously. Plasmid pY2 is functionally similar to pYESS 2.0 plasmid (Denard, Paresi et al. 2021), except that the protease is under the control of a β-estradiol (βE) inducible promoter, *(lexAbox)_3_-PminCYC1*, and the substrate cassette is driven by *pGal1.* Construction of the substrate cassette takes advantage of a *BsmBI* Golden Gate assembly whose success can be verified by green-white screening. The substrate cassettes were assembled by annealing phosphorylated oligonucleotides (oligos) flanked with the necessary overhangs that allows one to assemble these double-stranded parts in the desired order of FLAG, substrate, HA using a *BsmBI* Golden Gate assembly in a receiver plasmid, as described previously (Denard, Paresi et al. 2021). The enzyme can be cloned via a subsequent *BsaI* Golden Gate assembly. In addition, compared to the pYESS2.0 plasmid, unnecessary sequences were removed, making pY2 about 1000 base pair (bp) smaller than pYESS 2.0 (**Figure 1A**). Since we used the *(lexAbox)3-PminCYC1* for the first time, we characterized its strength compared to the *(lexAbox)4-PminCYC1* promoter in pYESS 2.0 and studied its leakiness using an mTurquoise fluorescent protein reporter. Although this promoter is weak, it does not show expression in the absence of inducer (**Supplemental Figure 1**) making it appropriate for our studies. Furthermore, a weak promoter is useful for this study because it allows us to quantify protease activities at low enzyme: substrate ratios. Thus, phenotypes observed can be generally translated to enzyme engineering and substrate profiling applications. We also constructed a plasmid, pY3, that is similar to pY2 except that the *(lexAbox)_3_-PminCYC1* and *pGal1* promoters were replaced with a bidirectional *pGal1-10* promoter. In this way, pY3 is like the original YESS plasmid but has the Golden Gate assembly features of pY2.

**Figure 1.**
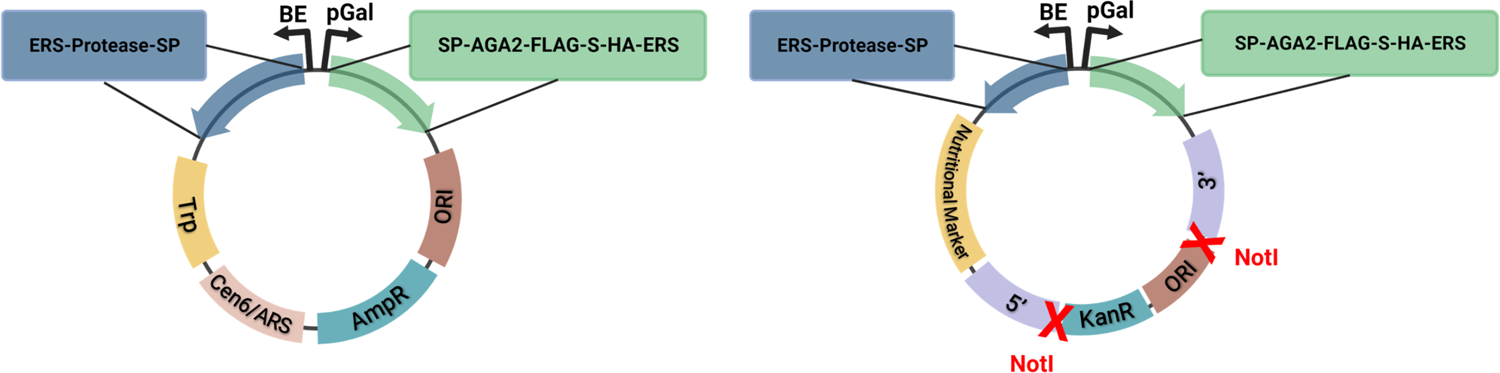
System components and expected display populations for plasmid and integration-based activity reporters. (A) Plasmid-based protease reporter system, inspired by pYESS, contains two functional cassettes – protease (blue: ER Retention Signal (ERS)-Protease-Signal Peptide (SP)) and substrate (green: SP-FLAG-Substrate (S)-HA-ERS) – with expression controlled via independent, inducible promoters. When transformed and expressed in yeast, the protease activity can be directly observed using flow cytometry of fluorescently-labeled substrate cassettes displayed on the cell surface. Expected displaying populations for plasmid-based systems will be broad (due to non-uniform expression) and include large non-displaying population of cells. (B) Integration-based protease reporter system contains the same functional cassettes as in (A), with the cassettes being flanked by chromosomal homology regions (5p and 3p). Linearization to expose homology regions occurs via NotI restriction enzyme.

To build integrative versions of pY2 and pY3, we chose two integration loci that exhibit high integration efficiencies, MET15 and LYS2 (Sadowski, Su et al. 2007). The homology arms were 500 to 800 bp 5’- and 3’-of the open reading frame and were amplified directly from LexA-hER-haB112-EBY100 (LLE) cells (**Table 1**) and cloned into a chloramphenicol backbone plasmid. We also chose one nutritional marker and one resistance marker, Leu2 and Hygromycin. Inspired by the plasmid construction blueprint from the yeast MoClo kit (Lee, DeLoache et al. 2015), we built two similar integrative plasmids; a MET15 integrative plasmid with a hygromycin resistance marker (MET-HYGRO) and a LYS2 integrative plasmid with a Leu2 nutritional marker (LYS-Leu2) (**Table 1**). The pY2 functional cassette was inserted into both integrative plasmids by removing the superfolder green fluorescent protein (sfGFP) with *BsaI* restriction enzyme digestion and HiFi DNA assembly (**Figure 1B**). A similar cloning scheme was followed to clone the pY3 functional cassette into our LYS-Leu2 integration plasmid (**Table 1) (Supplementary Figure 2)**.

**Table 1.**
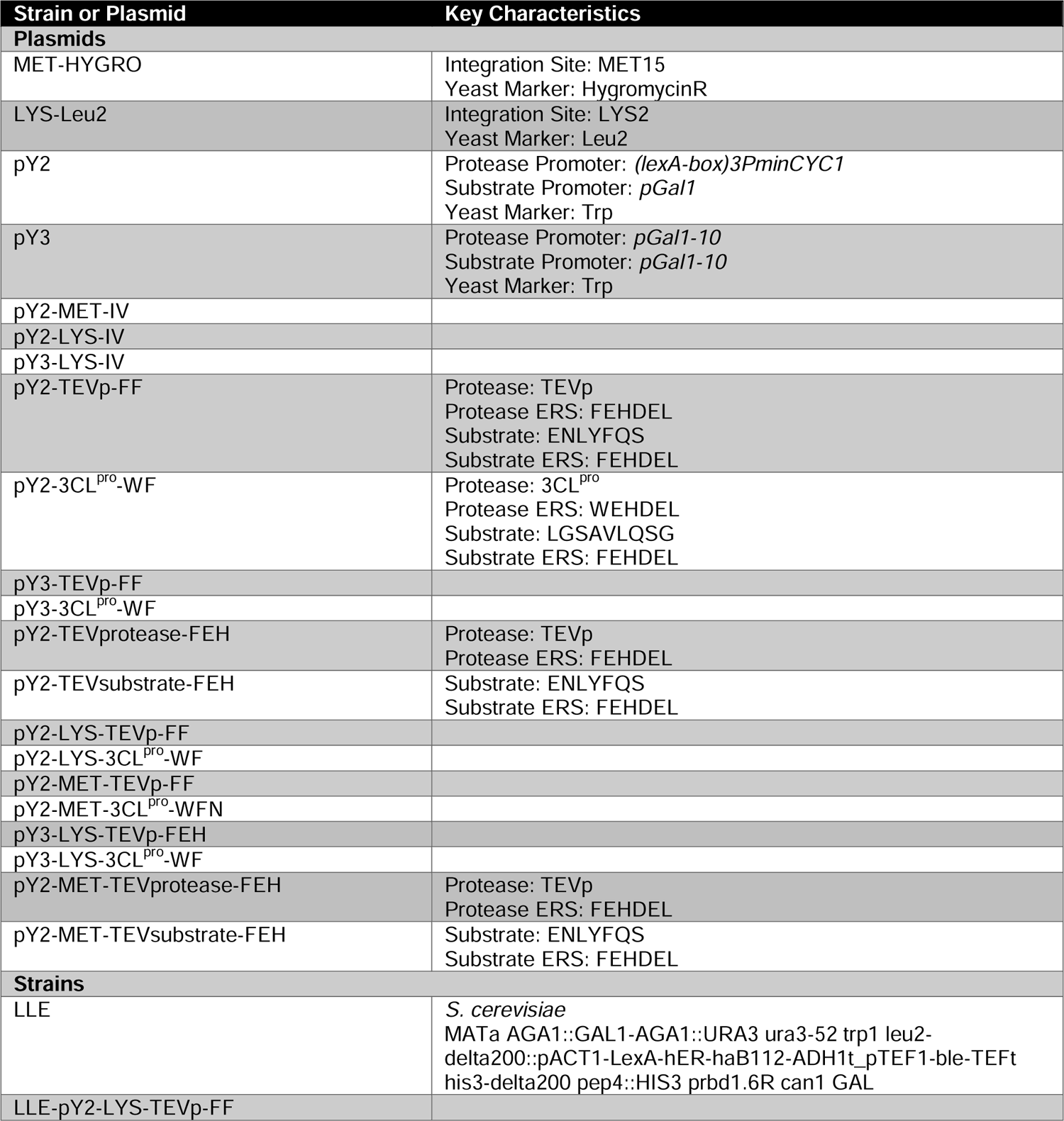
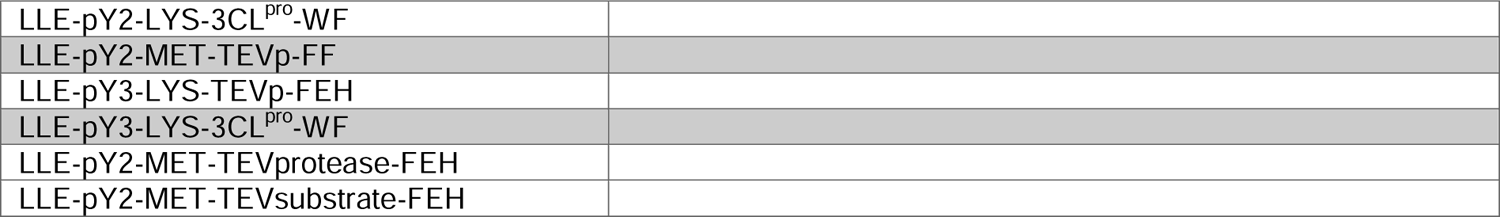
Plasmids and strains used in this study

### Integrative protease activity reporters enhance reliability and dynamic range of measurements on the yeast surface

When using a centromeric plasmid (CEN6/ARS) for cell surface display, about 10-30% cells do not show any display by flow cytometry (Boder and Wittrup 1997, Hackel and Wittrup 2008, Lee, DeLoache et al. 2015). Moreover, transcription noise is introduced due to plasmid copy number variation (4-8 copies per cell) (Jahn, Vorpahl et al. 2016, Shao, Rammohan et al. 2021, Fu, Patel et al. 2022). In our case, transcription noise is exacerbated because our plasmids carry two functional transcripts. As an example, we used pY2 to clone TEVp and its substrate under βE and galactose inducible promoters, respectively, with FEHDEL ERS on both cassettes (**Figure 2A, top plot)**. Here, we quantify protease activity by normalizing signal ratios (anti-FLAG PE/anti-HA Alexa 647) of the Protease On (blue) populations to the Protease Off (red) populations (see Methods). The two gene cassettes of our plasmid-based protease activity reporters result in a CV_HA_ and D_dB,HA_ of 1.01 and 4.16±0.67 dB, respectively. **(Supplementary Figure 3)**.

**Figure 2.**
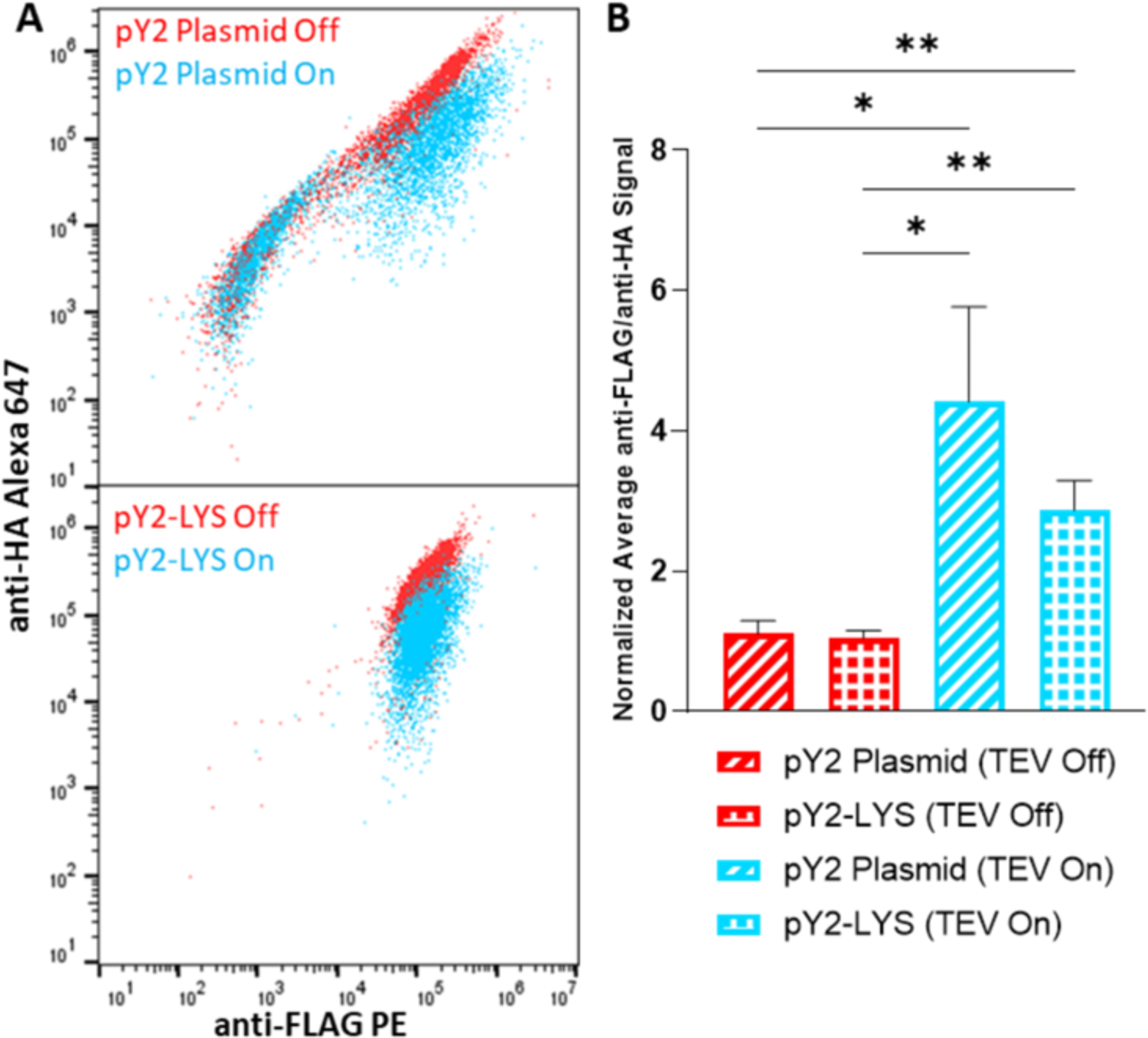
Using integration to improve display of TEVp activity in yeast. (A) Activity assay of tobacco etch virus protease (TEVp) conducted using flow cytometry to compare plasmid (top) and integrative (bottom) systems to confirm the use of integration to tighten expression control and display of cell populations with inactive TEVp represented in red and active TEVp under β-estradiol induction in blue. (B) Quantification of TEVp activity for both configurations, with a 4-fold and 2.7-fold increase in activity under βE for plasmid and integration, respectively. The activity of TEVp is quantified by a fold-change analysis of the anti-FLAG PE to anti-HA Alexa 647 mean fluorescence signal, normalized to the Protease Off (Red) ratio. Anti-FLAG PE/anti-HA Alexa 647 ≤1 correlates to protease inactivity, while >1 correlates to protease activity. No significant difference in activity was observed when comparing the substrate cassette present on a plasmid to the substrate cassette being integrated into the chromosome. Statistical significance between populations was determined by multiple unpaired t-tests. *p ≤0.05, **p ≤0.01, ***p ≤0.001, ****p ≤0.0001.

We reasoned that integrating the protease and substrate functional cassettes would alleviate this transcription noise and tighten the CV_HA_. We cloned TEVp and its substrate into pY2-LYS-IV (**Table 1**) using the streamlined cloning approach previously described (**Figure 1B**) and integrated the functional cassettes at the LYS2 integration site of our yeast strain. We assayed the cells using flow cytometry and directly compared expression variability and display of the two reporter systems. Integrating the functional cassettes resulted in a significant decrease in CV_HA_ (0.97), while maintaining a uniform D_dB,HA_ (3.87±0.92 dB), compared to the plasmid-based system **(Supplementary Figure 3)**. Analysis of protease activity levels illustrates that there are significant differences between Protease Off vs. Protease On in the integrative system. However, Protease On activity levels for the integrative system (4-fold increase in activity) do not differ significantly from the Protease On activity levels for the plasmid-based system (2.7-fold increase in activity) (**Figure 2B**). Overall, integration resulted in a tightly packed cell populations and the elimination of the non-displaying populations that are seen in the plasmid-based system (**Figure 2A, bottom plot)**. Interestingly, the mean protease activity remained the same in both systems, showing that integration has no impact on protease activity levels. While this is expected for non-toxic, sequence-specific proteases such as TEVp, the same may not hold true for promiscuous proteases where integration may result in both higher activity and more uniform active populations.

### pGAL1-10—driven transcription improves the performance of integrated TEVp

The galactose inducible promoters *pGal1* and *pGal1-10* are among the strongest promoters in *Saccharomyces cerevisiae (S. cerevisiae).* The original YESS system used a bidirectional *pGal1-10* promoter to drive expression of both enzyme and substrate cassettes. An integrative YESS with bidirectional *pGal1-10* would be highly beneficial to assay low-activity protease: substrate pairs (Mei, Zhai et al. 2017) and for drug screening (Taft, Georgeon et al. 2019). The βE and *pGAL1* promoters of pY2-LYS-IV were swapped with *pGAL1-10* to build pY3-LYS-IV (**Table 1**), which was integrated into our yeast strain and assayed for proteolytic activity using flow cytometry (see Methods). By expressing TEVp under *pGAL1-10*, we saw significant increase in activity (a 7.3-fold increase compared to 2.7-fold of pY2 configuration), keeping all other factors constant. As expected, CV_HA_ improved and D_dB,HA_ remained uniform for the pY3 integrative configuration compared to plasmid-based versions **(Supplementary Figure 5).**

### Expansion beyond the LYS2 integration site

In an effort to integrate multiple functional genes into the yeast chromosome we sought to expand beyond single site integration. Additionally, we were interested in confirming that nutritional markers and resistance genes have no influence on expression and subsequent protease activity. We integrated TEVp using pY2-MET-IV plasmid and compared the proteolytic activity to TEVp integrated using pY2-LYS-IV (**Table 1**). TEVp activity at MET15 was assayed using flow cytometry and quantified as described (see Methods). A small difference in fold change of activity when comparing integration sites (2.8 and 2.1-fold increase for LYS2 and MET15, respectively) was observed, providing confirmation that protease activity in yeast is largely independent of the integration loci chosen (**Figure 3B**). This conclusion was further confirmed with CV_HA_ and D_dB,HA_ calculations, with no significant differences observed between configurations **(Supplementary Figure 7)**.

**Figure 3.**
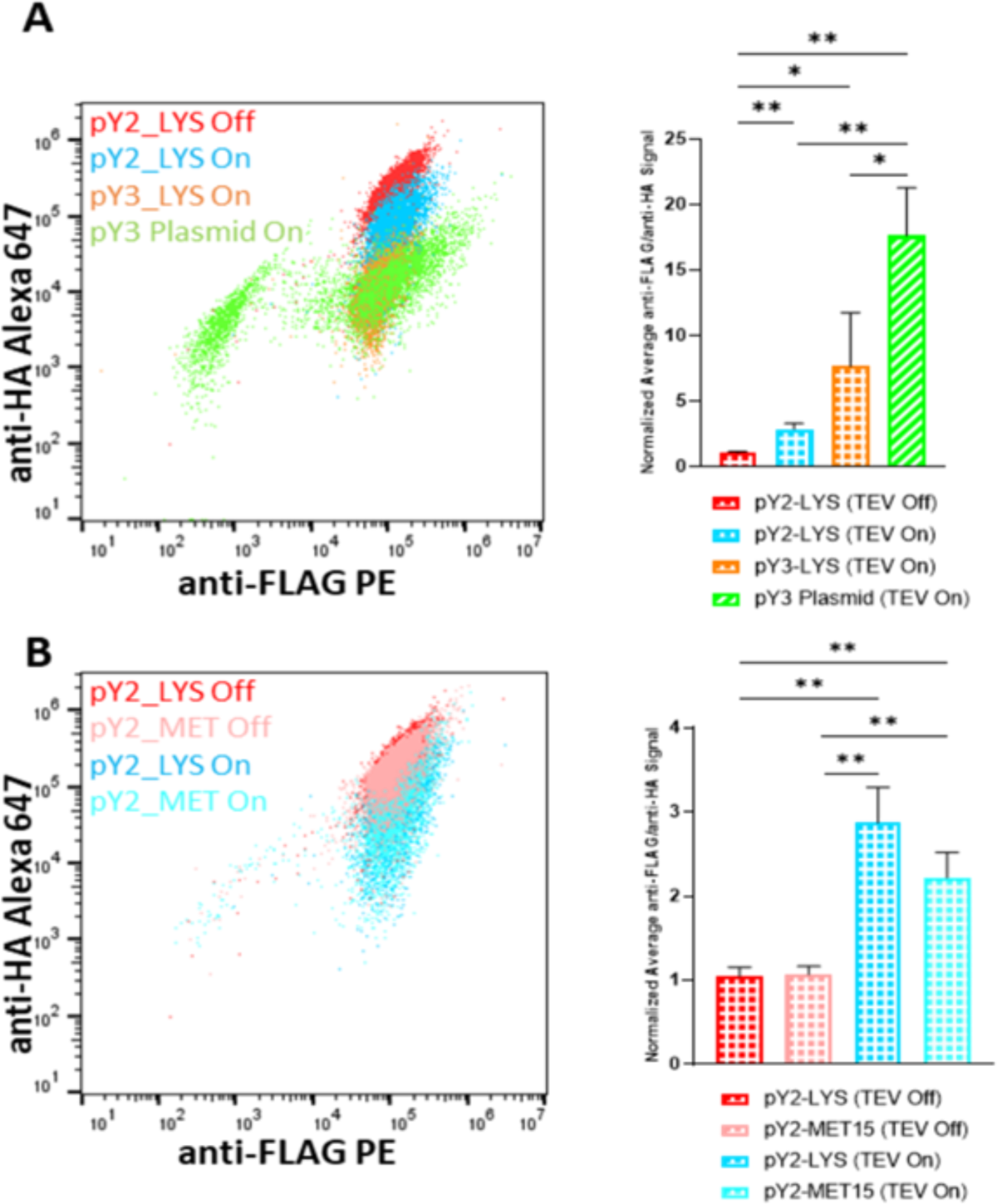
Expansion of platform to include multiple promoters and integration sites. (A) The activity of TEVp integrated at LYS represented by three cell populations: inactive protease (red), active protease under βE control (pY2, blue) corresponding to a 2.7-fold increase in activity, and active protease under galactose control (pY3, orange) corresponding to a 7.3-fold increase in activity. (B) Representation that integrating TEVp at two different integration sites, LYS2 and MET15, does not result in changes in observed activity. Inactive protease populations represented by red (LYS2) and pink (MET15) and active protease populations represented by blue (LYS2) and turquoise (MET15) – no significant difference in fold-change activities in both cases. Statistical significance between populations was determined by multiple unpaired t-tests. *p ≤0.05, **p ≤0.01, ***p ≤0.001, ****p ≤0.0001.

### Quantifying disease-relevant protease activity using integrative reporter platform

The proteolytic activity of TEVp has been well documented in yeast (Yi, Gebhard et al. 2013, Cesaratto, López-Requena et al. 2015, Sanchez and Ting 2020, Denard, Paresi et al. 2021, Dyer and Weiss 2022), making it a suitable choice for our initial protease of interest. However, it is necessary to expand the system to study proteases of different activity ranges and interests in human health. 3-chymotrypsin like protease(3Cl^pro^) the main protease involved in severe acute respiratory syndrome coronavirus-2 (SARS-CoV-2) replication, was picked due to its distinct canonical substrate (LGSAVLQSG) (Chen, Yiu et al. 2020), along with the increased interest in inhibiting 3CL^pro^ activity for therapeutic benefit (Koulgi, Jani et al. 2021, Owen, Allerton et al. 2021, Rajpoot, Alagumuthu et al. 2021). 3CL^pro^ was cloned into both plasmid and integration versions of our βE-driven protease reporters and assayed as described (see Methods) (**Figure 4A**). As observed with TEVp, we report that the fold change increases in activity when 3CL^pro^ expression was induced were not significantly different between the plasmid and integrative systems (2.5 and 2.1-fold increase in activity, respectively). This trend was also consistent when 3CL^pro^ was under the stronger galactose promoter (14 and 12.4-fold increase in activity across plasmid and integrative systems, respectively) (**Figure 4B**). Tighter populations and stable D_dB,HA_ for integrated 3CL^pro^, correlating to uniform transcription and expression of gene cassettes, are also reported **(Supplementary Figure 9)**.

**Figure 4.**
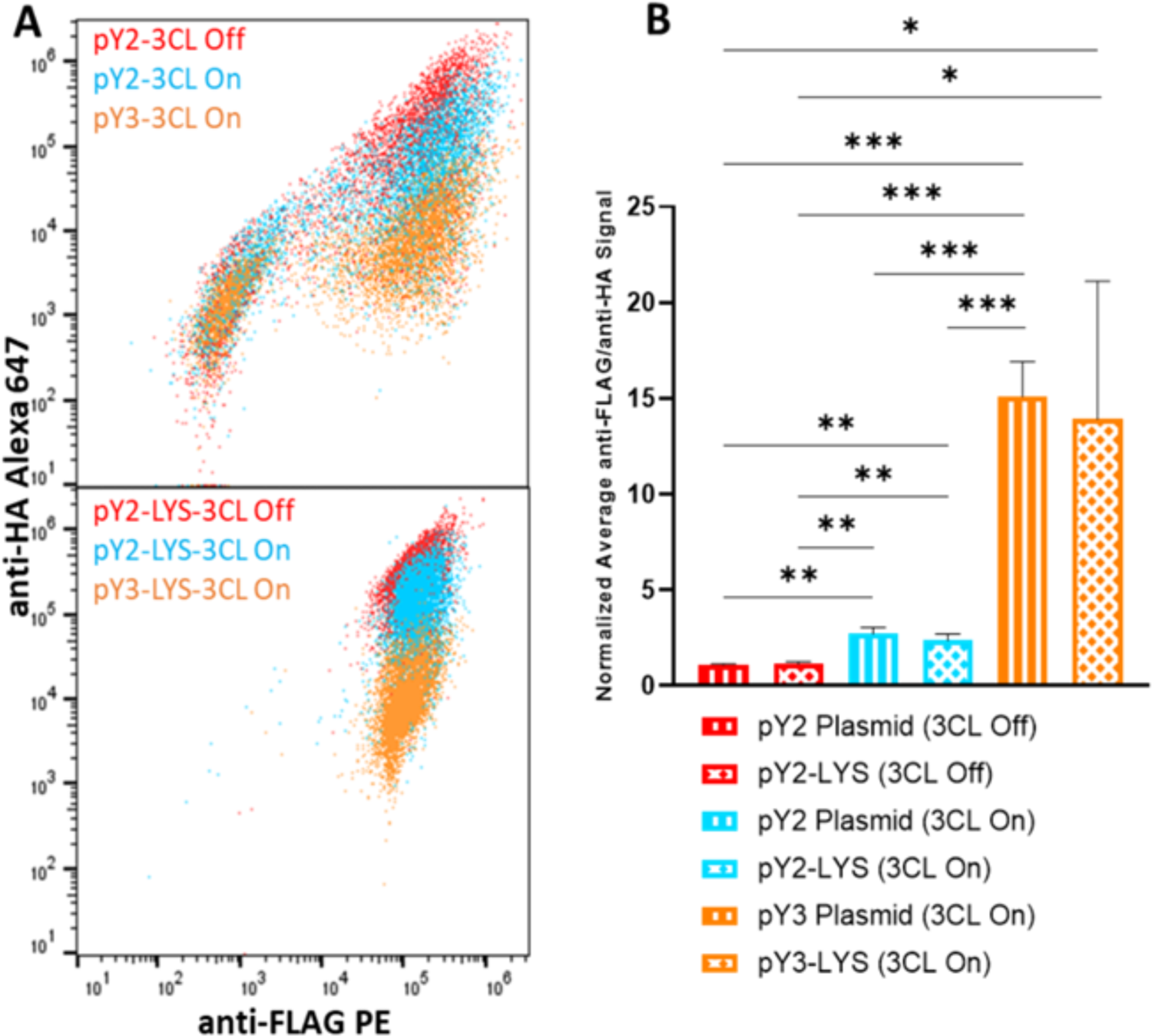
Analysis of disease-relevant proteases using SIVs. (A) Activity assay of 3CL (SARS-CoV-2 main protease) conducted using flow cytometry to compare plasmid (top) and integrative (bottom) systems to confirm the use of integration to tighten expression control and display of cell populations with inactive 3CL represented in red, active 3CL under βE induction in blue and active 3CL under galactose induction in orange. (B) Quantification of 3CL activity for all configurations, with a 2.5-fold and 2.1-fold increase in activity under βE(Blue) for plasmid and integration, respectively, and a 14-fold and 12.4-fold increase in activity under galactose induction (Orange). The activity of TEVp under BE control is quantified by a fold-change analysis of the anti-FLAG PE to anti-HA Alexa 647 mean fluorescence signal, normalized to the Protease Off (Red) ratio. Anti-FLAG PE/anti-HA Alexa 647 ≤1 correlates to protease inactivity, while >1 correlates to protease activity. No significant difference in activity was observed when comparing the substrate cassette present on a plasmid to the substrate cassette being integrated into the chromosome. Statistical significance between populations was determined by multiple unpaired t-tests. *p ≤0.05, **p ≤0.01, ***p ≤0.001, ****p ≤0.0001.

### Integration streamlines and enhances biochemical studies performed in the yeast ER

YESS allows one to perform several high-throughput investigations, including enzyme engineering, substrate profiling, deep mutational scanning, and drug screening. For protease engineering and deep mutational scanning campaigns, the substrate cassette remains constant. Conversely, for substrate profiling, the protease remains constant. Lastly, for drug screening, both substrate and enzyme can be chromosomally integrated. Chromosomal integration of the constant transcriptional cassette eliminates unwanted mutations and minimizes plasmid size and transcriptional noise. To enable these various assays, we built split-cassette systems in which one functional cassette is integrated into the chromosome, and the other remains on a plasmid transformed into the integrative strain. We established two yeast strains, one with our LE-driven TEVp cassette integrated and the other with the galactose-inducible TEV substrate cassette integrated, both at MET15. Into these strains, we transformed the counterpart cassette plasmid such that each contained a protease and a substrate and assayed them using flow cytometry (see Methods) (**Figure 5A and 5B**). The integrated substrate cassette coupled with the plasmid-based protease cassette produced a linear cell population, beginning in the non-displaying region (low anti-FLAG and anti-HA signals) and travels diagonally upward to the region associated with high levels of both signals **(Supplementary Figure 11A)**. This linearity was also observed for the strain containing TEVp integrated into the chromosome **(Supplementary Figure 11B)**. Additionally, we observed that the TEVp integration system resulted in higher levels of activity than the TEVp plasmid-based system (5.9 and 3.3-fold increase, respectively) (**Figure 5C**). We hypothesize that this can be attributed to the potential of non-complete transcription of the protease gene when it is present on a plasmid, compared to complete transcription when integrated into the chromosome. This split-cassette platform can be directly translated to library screening and expands the experimental applications of this integrative system.

**Figure 5.**
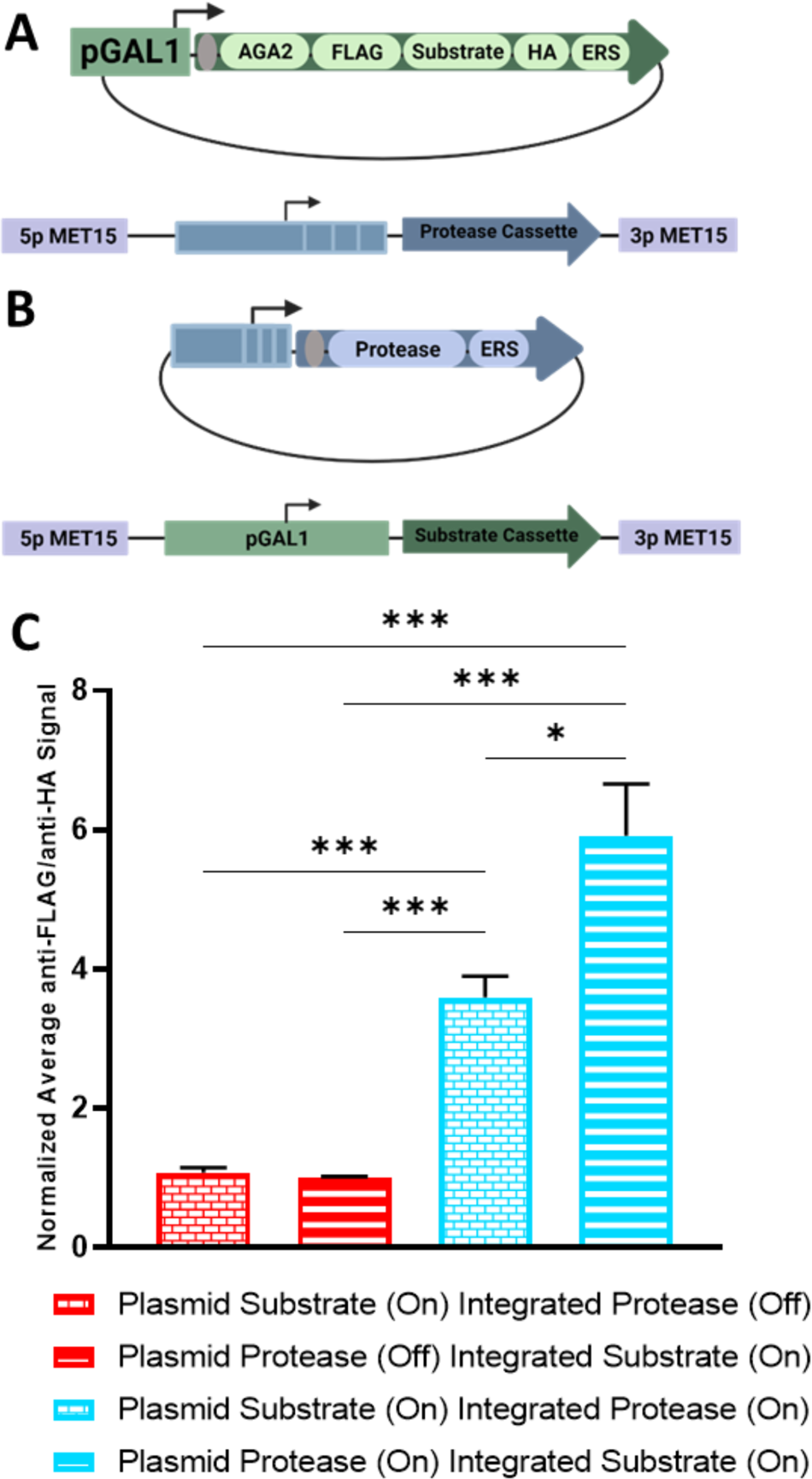
Integration strategy for library screening. (A) Schematic of plasmid-integration system in which the substrate cassette (under galactose induction) is present on a plasmid and the protease cassette (under β-estradiol induction) is integrated into the yeast chromosome at MET17. (B) Schematic of plasmid-integration system in which the protease cassette (under β-estradiol induction) is present on a plasmid and the substrate cassette (under galactose induction) is integrated into the yeast chromosome at MET17. (C) Quantification of protease activity for both configurations, with 3.3-fold and 5.9-fold increases in the activity being observed for plasmid and integration of substrate cassette, respectively. The activity of TEVp under BE control is quantified by a fold-change analysis of the anti-FLAG PE to anti-HA Alexa 647 mean fluorescence signal, normalized to the Protease Off (Red) ratio. Anti-FLAG PE/anti-HA Alexa 647 ≤1 correlates to protease inactivity, while >1 correlates to protease activity. No significant difference in activity was observed when comparing the substrate cassette present on a plasmid to the substrate cassette being integrated into the chromosome. Statistical significance between populations was determined by multiple unpaired t-tests. *p ≤0.05, **p ≤0.01, ***p ≤0.001, ****p ≤0.0001.

## Conclusion

The YESS system has been widely used within the community to study PTM-enzymes and their substrates to enable engineering and modulating of enzyme specificities and selectivities. To increase the HT capability of this system for assaying and characterizing enzyme-substrate interactions, we built and characterized integrative YESS plasmids pY2-LYS-IV, pY2-MET-IV, and pY3-LYS-IV (**Table 1**). These vectors provide a streamlined cloning and transformation approach to insert enzyme activity reporters into the chromosome of yeast for yeast surface display (YSD)-based assays of proteolytic activity using flow cytometry. This integrative system can efficiently quantify TEVp activity on its canonical substrate with tight expression control using chromosomal integration, while maintaining comparable D_dB,HA_ across both system configurations. This was also seen with 3CL^pro^, a main protease involved in SARS-CoV-2 viral replication. To provide a system that can be used to screen multiple enzyme-substrate reporters concurrently, we provided validation that TEVp activity was not hindered through use of multiple integration sites and different selection markers. Use of this system for evolutionary engineering of PTM-enzymes is achieved through our split-integration system in which we show that the constant (non-engineered) cassette can be integrated solely into the chromosome while the variable (engineered) cassette can be transformed and retained on a plasmid. Overall, we provide the community with a streamlined approach for screening and characterizing enzyme-substrate interactions through chromosomal integration of activity reporters to expand the reach of yeast-based enzyme engineering.

## Methods and Materials

### Plasmid construction

#### Protease and substrate preparation

To assemble the protease cassette the 3CL^pro^ was amplified from the plasmid pLEX307-SARS-CoV-2-3CL, a gift from Alejandro Chavez (Addgene plasmid # 160278; http://n2t.net/addgene:160278; RRID: Addgene 160278). In addition, the TEV protease was amplified from the plasmid pRK793, a gift from David Waugh (Addgene plasmid # 8827; http://n2t.net/addgene:8827; RRID: Addgene_8827). The desired ERS strength was encoded onto the primers used to amplify the protease genes. The resulting polymerase chain reaction (PCR) products were then purified (ZymoResearch #D4003) and prepared for subsequent insertion into the receiver plasmids. To prepare the TEV substrate two oligos were annealed using T4 Polynucleotide Kinase (NEB#M0201S) and T4 DNA ligase buffer (NEB#B0202S). This same annealing method was used to make the 3CL substrate, FLAG, HA and substrate ERS parts. **Supplemental Table 1** contains oligo sequences used for protease and substrate preparation.

#### Plasmid-based system

pY2 plasmid was constructed via Gibson Assembly (Gibson, Young et al. 2009) of a CEN6-TRP backbone and the protease and substrate cassette configuration that defines pY2. Each cassette contains unique cut sites, allowing for streamlined cloning at both loci to functionalize the plasmid. To clone into the substrate cassette, a *BsmBI* Golden Gate (Engler, Kandzia et al. 2008) reaction was performed to remove sfGFP and replace it with the cassette components (P1-FLAG, P2-Substrate, P3-HA, P4-ERS). This plasmid was transformed into competent *Escherichia coli (E. coli)* (NEB#C2984H), purified (QIAGEN #27106) and sent for sequencing to confirm correct assembly (Plasmidsaurus, standard plasmid). Once confirmed, a *BsaI* Golden Gate reaction (Engler, Kandzia et al. 2008) was done to replace the multiple-cloning site with the protease sequence (containing an ERS). Amplification and sequencing of this plasmid was done as previously described. Similar construction was done for the pY3 plasmid replacing the βE promoter and the *pGal1* promoter with a *pGal1-10* promoter. **Supplementary Table 2** contains thermocycler protocols for the reactions mentioned.

For enzyme activity assays, protease-substrate containing plasmids were transformed into an engineered *S. cerevisiae* strain (EBY100) expressing a LexA-hER-haB112 transcription factor (necessary for LE induction). Transformation protocol followed the Frozen-EZ Yeast Transformation II Kit (ZymoResearch #T2001), with transformed cells plated on yest nitrogen base (YNB) casamino acid (CAA) 2% Glucose and grown at 30°C for 2-3 days. Mature colonies were picked and grown in 1 mL YNB CAA 2% Glucose 2% Raffinose media at 30°C for 16-24 hours, until saturated.

#### Integration-based system

A PCR was performed on genomic DNA from *S. Cerevisiae* (ZymoResearch#D2002) and then purified (Zymo#D4004) to obtain the 5’ and 3’ homologies for the respective integration sites and overhangs. A *BsaI* Golden Gate **(Supplementary Table 2)** using the MoClo Yeast Toolkit (Lee, DeLoache et al. 2015) was then done with the homologies to construct the initial green fluorescent protein (GFP) plasmid and then transformed into competent *E.coli* (NEB#C2984H) and subsequently sequenced (Plasmidsaurus, standard plasmid). These initial plasmids contained unique cut sites flanking the sfGFP for efficient cloning. The pY2 and pY3 functional cassettes were then amplified off the previously built pY2 and pY3 plasmids and inserted into the integration vectors via Gibson Assembly **(Supplementary Table 2)**, replacing sfGFP with the empty protease and substrate cassettes. Amplification and sequencing of these plasmids was done as previously described. Once sequencing was confirmed substrates and proteases could be inserted using *BsmBI* and *BsaI* Golden Gate reactions, respectively **(Supplementary Table 2)**.

To integrate the protease-substrate cassettes, integrative vectors were linearized via a *NotI* (NEB #R3189L) digestion of 3-5 ug of plasmid. The digestion was run at 37°C for 5 hours and gel purified (ZymoResearch #D4001) to capture the top band (approximately 6 kbp). Linear cassettes were transformed into an engineered *S. cerevisiae* strain (EBY100) expressing a LexA-hER-haB112 transcription factor (necessary for LE induction). Transformation for integration followed the Lithium Acetate (LiAc) transformation protocol outlined by Gietz and Schiestl (Gietz and Schiestl 2007), with an added 2 hour outgrow in YPD at 30°C following the 42-minute incubation at 42°C for additional recovery. Transformations were plated on selection media, YPD Hygromycin (MET15 integration site) and SC-LEU (LYS2 integration site) and grown at 30°C for 2-4 days. Mature colonies were picked and grown in 1 mL selection media at 30°C for 16-24 hours, until saturated.

### Building pY2-TEVprotease-FEH and pY2-TEVsubstrate-FEH

A Q5 mutagenesis reaction (NEB#M0491) was used to remove the protease or substrate cassettes from the pY2-MET-TEVp-FF plasmid (**Table 1**). Once the Q5 reaction was complete, a KLD reaction (NEB#M0554S) was done to reconnect the plasmid, creating pY2-MET-TEVsubstrate-FEH and pY2-MET-TEVprotease-FEH plasmids that contain only TEV substrate or TEVp, respectively (**Table 1**). These plasmids were then transformed into competent *E. coli* (NEB#C2984H), purified (QIAGEN #27106) and sent for sequencing to confirm correct assembly (Plasmidsaurus, standard plasmid). The same protocol was starting with pY2-TEVp-FF to obtain both pY2-TEVsubstrate-FEH and pY2-TEVprotease-FEH plasmids (**Table 1**). Once sequencing of these plasmids was confirmed (plasmidsaurus), the integration versions were transformed separately via the LiAc cell transformation protocol described. Competent LLE cells were then created using the Zymo Research Frozen-EZ Yeast Transformation II Kit (Zymo#T2001). When the component cells containing either Integrated TEVsubstrate or TEVp were complete, the plasmid counterpart was then transformed using the plasmid transformation protocol described above. Supplemental Table 5.2 contains oligo sequences used for Q5 mutagenesis PCR.

### Protease activity assays using flow cytometry

#### Cell Preparation

Saturated cultures (24-hour inoculation from colony, 1 mL media, 30°C, 250 RPM) were collected, and the optical density (OD_600_) was taken to determine the approximate amount of saturated culture required to outgrow the cells starting from an OD_600_ of 1 in 0.25 mL media. In a deep well 96-well plate (ThermoFisher Scientific, 260252), cultures were inoculated to desired outgrow OD_600_ in 0.25 mL YNB CAA 2% Glucose, 2% Raffinose and grown to an OD_600_ between 2-4 (5-hour outgrow, 30°C, 800 RPM on a plate shaker). OD_600_ of outgrown cultures was calculated, and the approximate amount of culture required to induce at an OD_600_ of 0.5 in 0.25 mL media was determined. Required culture volumes, two per sample (one for Protease Off and one for Protease On), were washed in minimal galactose-rice media (YNB CAA 2% Galactose) to wash residual glucose in a second deep well 96-well plate. Once the wash supernatant was removed, cells were induced for protein expression. Protease Off samples were induced with 0.25 mL YNB CAA 2% Galactose for substrate expression. Protease On samples were induced with 0.25 mL YNB CAA 2% Galactose and βE (at 2 uM) for substrate and protease expression. The induction plate was grown at 30°C for 12-16 hours, shaking at 800 RPM. OD_600_ of induced cultures was taken and used to determine approximate volumes of culture to stain two million cells with fluorescently labeled antibodies (anti-FLAG PE, Biolegend, cat# 637309, anti-HA Alexa 647, Biolegend, cat #682404). Cells were washed in PBS 0.5% BSA (0.5% BSA, Goldbio 9048-46-8) in a round bottom 96-well plate (ThermoFisher Scientific, 174929) and stained (for 2 million cells: 0.5 uL anti-FLAG PE and 1 uL anti-HA Alexa 647) at RT for 90 minutes. Stained cells were washed with PBS 0.5% BSA and resuspended for cytometric assay (NL Cytek 3000, Cytek Biosciences).

#### Cytometric Assay

Following established assay protocols using flow cytometry, stained cells were analyzed for fluorescence in the B4-A, correlating with high display levels of our FLAG epitope tag (stained with anti-FLAG PE) and R2-A, correlating with high display levels of our HA epitope tag (stained with anti-HA Alexa 647) channels of the Nb Cytek 3000. Initial yeast cell populations were captured using a forward scatter (FSC-A) (x) vs. side scatter (SSC-A) (y) plot in log scale using a square gate at ≥10^4^ FSC-A and ≥10^4^ SSC-A **(Supplementary Figure 12)**. These cells were then displayed on a B4-A (x) R2-A (y) plot and displaying cells were captured using a straight gate that captured all cells at ≥10^4^ B4-A display and all ranges of R2-A display **(Supplementary Figure 13)**. FSC Files were obtained and further analyzed (see *Quantification of protease activity*) using FlowJo (Version 10.8.0).

### Quantification of protease activity

The substrate cassette is designed so that proteolytic cleavage can be directly related to the presence of one or both epitope tags that are flanking substrate sequences when bound to the yeast surface. If a protease was induced (Protease On), the substrate is expected to be cleaved prior to surface display, resulting in a loss of the anti-HA fluorescence signal and a retention of the anti-FLAG fluorescence signal that remains present in the cassette. If a protease is not induced (Protease Off), the displayed substrate cassette is expected to retain anti-HA and anti-FLAG fluorescent signals. Due to this design, one can establish a ratio of anti-FLAG/anti-HA average fluorescent signals obtained using flow cytometry to create a fold-change quantification of protease activity related to substrate cleavage. Using the Protease Off population as the baseline, anti-FLAG/anti-HA ratios of each colony in the triplicate are normalized and averaged to the first colony and then averaged to establish a value of approximately one. The Protease On population ratios are subsequently normalized to their Protease Off equivalent and averaged. Using this framework, one can define an active protease by a normalized fold change ratio >1 compared to the Protease Off baseline. These values can be used to directly compare configurations of the functional cassettes in our system of a single protease while allowing for cross-comparison of different proteases assayed.

Additional quantification more directly related to the system’s configuration (plasmid versus integration) can be achieved via comparing population spreads obtained in cytometric assays. The population spread can be directly calculated using taking the CV_HA_ of the anti-HA Alexa 647 signal of displaying cell population in FlowJo (Version 10.8.0). CV_HA_ can be translated to the transcription variability within the cells and is represented as a percentage, with lower percentages indicating tighter uniformity of protein transcription. To determine activity levels within a population, we used Equation 1 to calculate D_dB,HA_, the absolute value of the difference in amplitude of the mean fluorescence of anti-HA signal (R2-A) between Protease On and Off populations. All reported D_dB,HA_ values are representative of the average D_dB,HA_ across each triplicate. D_dB,HA_ is representative of protease activity on target substrate, with a larger magnitude corresponding with increased activity.

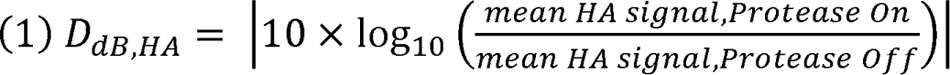

All quantification measurements are analyzed using a multiple t test with p values corresponding to the following scale: *p ≤0.05, **p ≤0.01, ***p ≤0.001, ****p ≤0.0001.

It is noted that the proteases presented in this paper were all determined to be active in the yeast-based system, however there is the potential that certain human and disease-relevant proteases may be natively inactive in yeast.

