## Supplemental figures for "Modular and integrative activity reporters enhance biochemical studies in the yeast ER"

### Supplemental Information

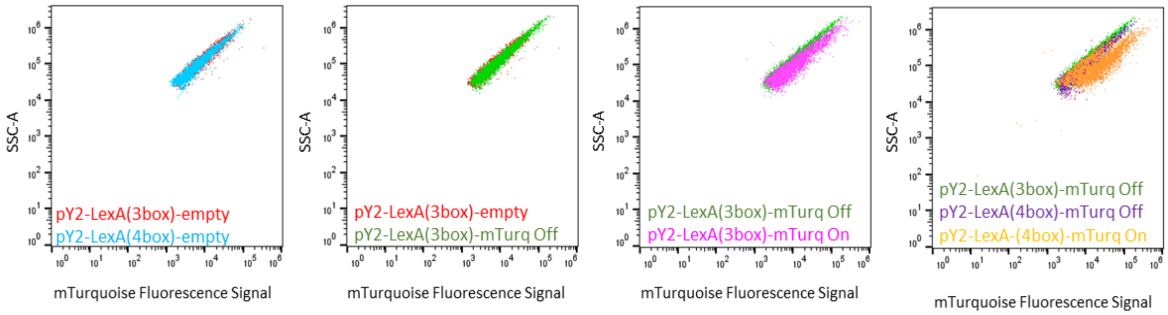


**Supplementary Figure 1. Dot plot representation of LexA promoter configuration influence on expression.** To explore the relationship between promoter configuration and expression, two mTurquoise2 (mTurq) plasmids were created, one containing a 3-box LexA promoter and the other, 4-box LexA promoter. Depicted here are comparisons of (A) 3box and 4box configurations expressing an empty gene cassette, (B) 3box configuration expressing an empty cassette (red) and an uninduced mTurquoise2 (mTurq) cassette under a 3box promoter (green), (C) 3box configuration expressing mTurq uninduced (green) and induced (pink), and (D) uninduced 3box and 4box configurations driving mTurq expression (green and purple, respectively) compared to induced 4box-driven expression of mTurq (orange).

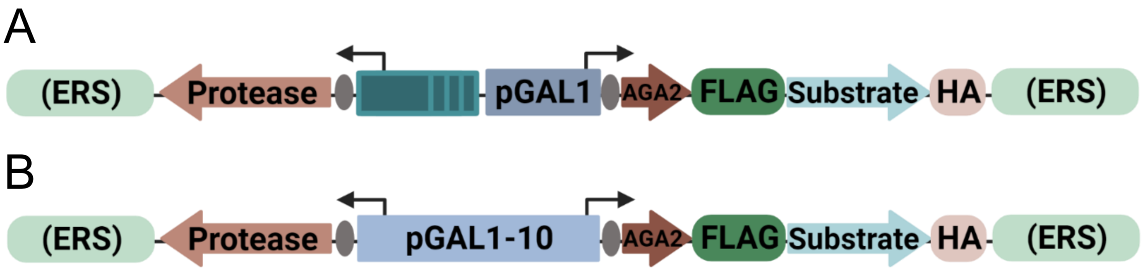


**Supplementary Figure 2. Schematic of functional cassettes.** (A) pY2 functional cassettes, protease cassette expression driven by *(lexA-box)3PminCYC1* and substrate cassette expression driven by *pGal1*. (B) pY2 functional cassettes, with both cassettes driven by bi-directional *pGal1-10.*

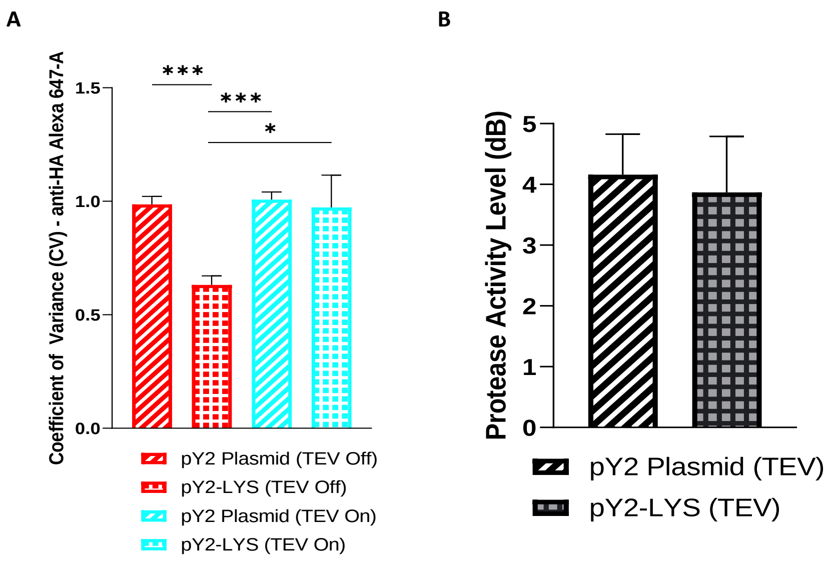


**Supplementary Figure 3. Population spread and signal-to-noise ratio improved through integration of TEVp.** (A) Coefficient of Variance ($CV_{HA}$) of the two system configurations calculated on the displaying yeast cells and the variability of anti-HA Alexa 647-A fluorescence signal across the cell populations. $CV_{HA}$, a quantification of population spread, is dependent on the uniform expression of the genes that translate to a signal being observed. (B) Protease activity level ($D_{dB,HA}$) calculated using Equation 1 and represented in units of dB. No significance was observed between activity levels observed in both plasmid and integration-based systems. Statistical significance between populations was determined by multiple unpaired t-tests. *p ≤0.05, **p ≤0.01, ***p ≤0.001, ****p ≤0.0001.

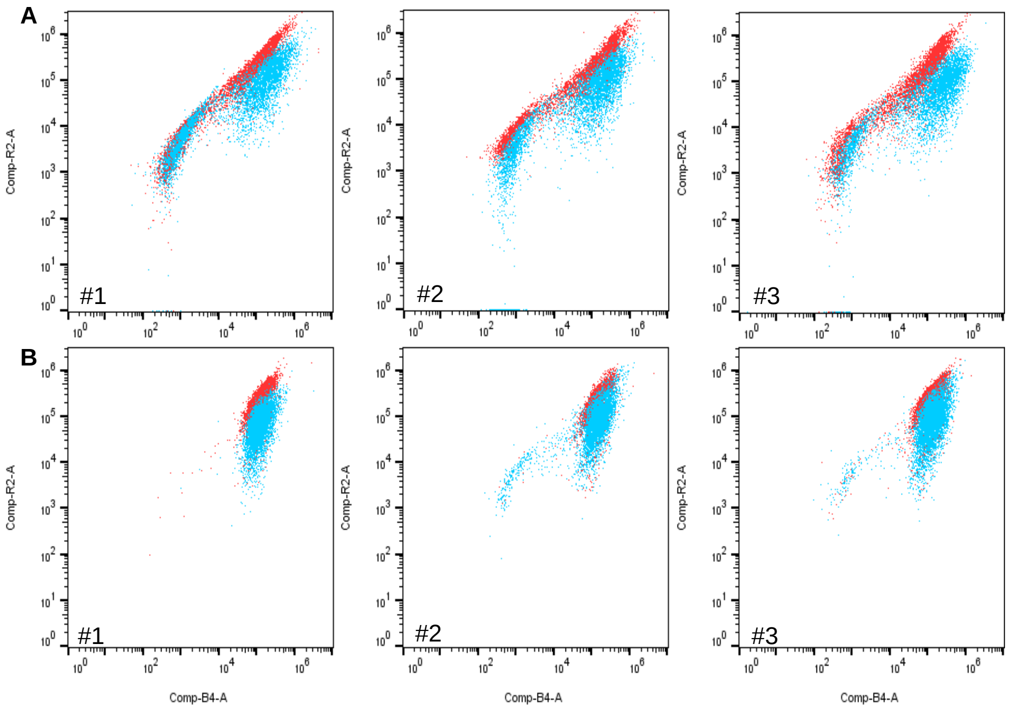


**Supplementary Figure 4. Triplicates of TEVp activity assay.** (A) TEVp in plasmid-based system, pY2 configuration. TEV Off (red) and TEV On (blue) populations presented on anti-FLAG PE (B4-A, x-axis) v. anti-HA Alexa 647 (R2-A, y-axis) dot plots. Populations presented in log scale. (B) TEVp in integration-based system, pY2 configuration integrated at LYS2 site. TEV Off (red) and TEV On (blue) populations presented on anti-FLAG PE (B4-A, x-axis) v. anti-HA Alexa 647 (R2-A, y-axis) dot plots. Populations presented in log scale.

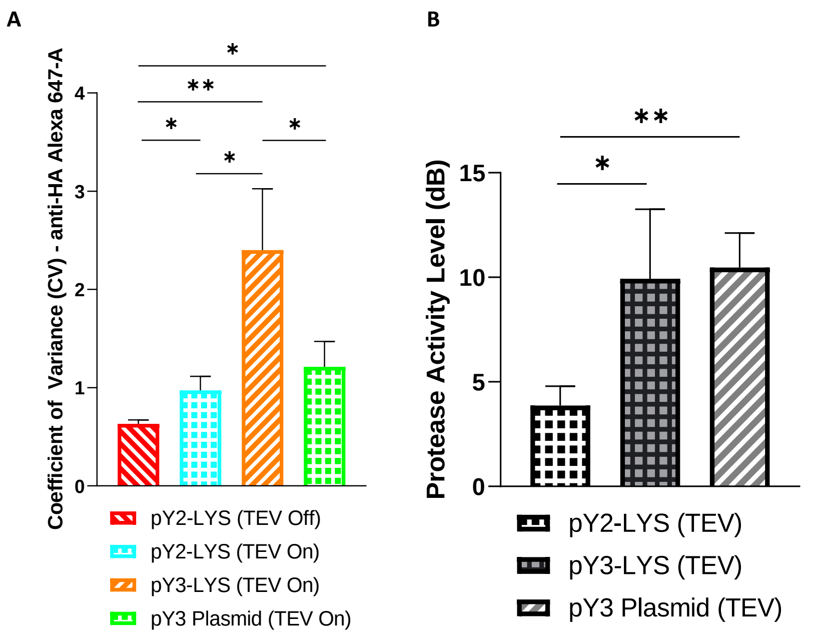


**Supplementary Figure 5. Population spread and signal-to-noise ratio are independent of activity level of TEVp.** (A) Coefficient of Variance ($CV_{HA}$) was calculated on the displaying yeast cells and the variability of anti-HA Alexa 647-A fluorescence signal across the cell populations. $CV_{HA}$, a quantification of population spread, is dependent on the uniform expression of the genes that translate to a signal being observed but is independent of activity level. Activity level of TEVp in this system is dependent on promoter selection. Comparison represented depicts the difference in $CV_{HA}$ observed across pY2 and pY3 plasmid versus integration configurations of TEVp, in addition to pY3 plasmid (green). (B) Protease activity level ($D_{dB,HA}$) calculated using Equation 1 and represented in units of dB. No significance was observed between activity levels observed in both plasmid and integration-based systems when protease expression was under the same promoter (pY3, *pGAL1-10*). Statistical significance between populations was determined by multiple unpaired t-tests. *p ≤0.05, **p ≤0.01, ***p ≤0.001, ****p ≤0.0001.

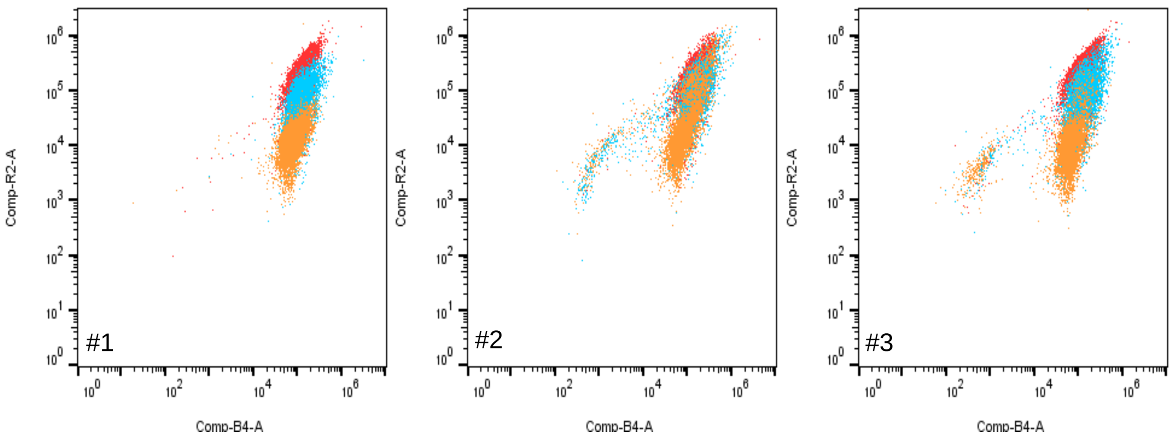


**Supplementary Figure 6. Triplicates of TEVp promoter swap activity assay.** TEVp in integration-based system, integrated at LYS2 site. TEV Off (red), pY2 TEV On (blue) and pY3 TEV On (orange) populations presented on anti-FLAG PE (B4-A, x-axis) v. anti-HA Alexa 647 (R2-A, y-axis) dot plots. Populations presented in log scale.

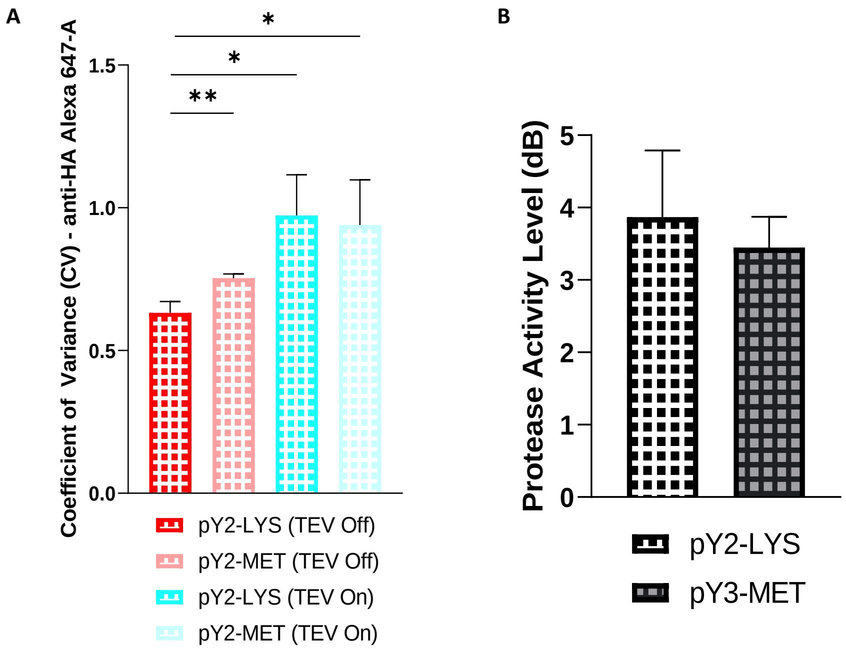


**Supplementary Figure 7. Population spread and signal-to-noise ratio are independent of integration site.** (A) Coefficient of Variance ($CV_{HA}$) of the of the two integration configurations calculated on the displaying yeast cells and the variability of anti-HA Alexa 647-A fluorescence signal across the cell populations. $CV_{HA}$, a quantification of population spread, is dependent on the uniform expression of the genes that translate to a signal being observed. (B) Protease activity level ($D_{dB,HA}$) calculated using Equation 1 and represented in units of dB. No significance was observed between activity levels observed in both integration-based systems. Statistical significance between populations was determined by multiple unpaired t-tests. *p ≤0.05, **p ≤0.01, ***p ≤0.001, ****p ≤0.0001.

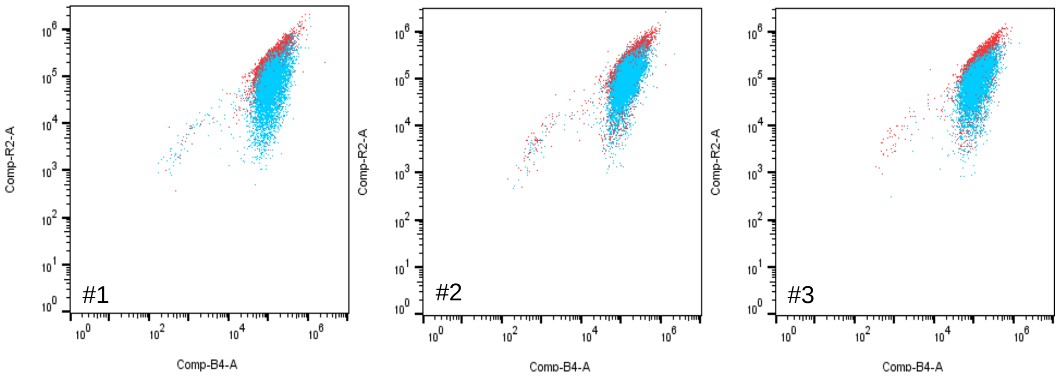


**Supplementary Figure 8. Triplicates of TEVp integration site swap activity assay.** TEVp in integration-based system, pY2 configuration integrated at MET15 site. TEV Off (red) and TEV On (blue) populations presented on anti-FLAG PE (B4-A, x-axis) v. anti-HA Alexa 647 (R2-A, y-axis) dot plots. Populations presented in log scale.

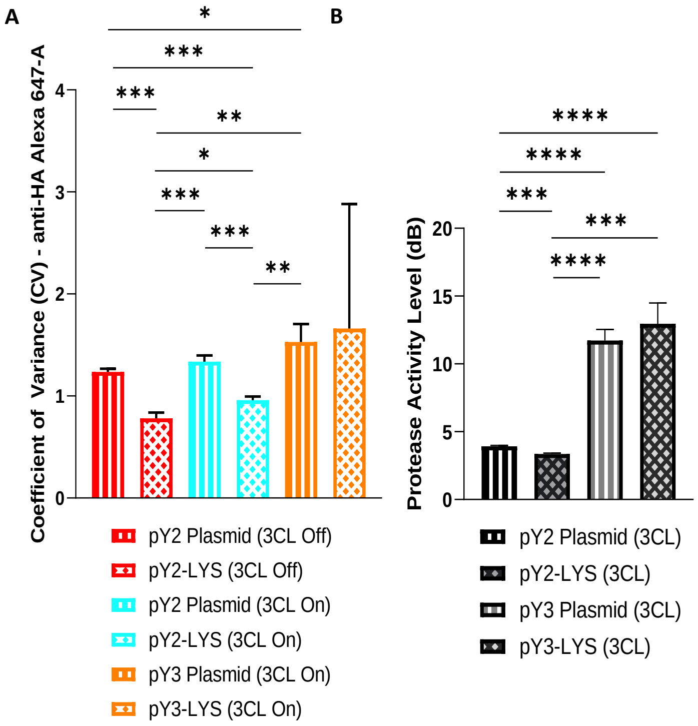


**Supplementary Figure 9. Population spread and signal-to-noise ratio improved through integration of 3CL$^{pro}$.** (A) Coefficient of Variance ($CV_{HA}$) of the of the two system configurations, in addition to two promoter configurations, was calculated on the displaying yeast cells and the variability of anti-HA Alexa 647-A fluorescence signal across the cell populations. $CV_{HA}$, a quantification of population spread, is dependent on the uniform expression of the genes that translate to a signal being observed. (B) Protease activity level ($D_{dB,HA}$) calculated using Equation 1 and represented in units of dB. Statistical significance between populations was determined by multiple unpaired t-tests. *p ≤0.05, **p ≤0.01, ***p ≤0.001, ****p ≤0.0001.

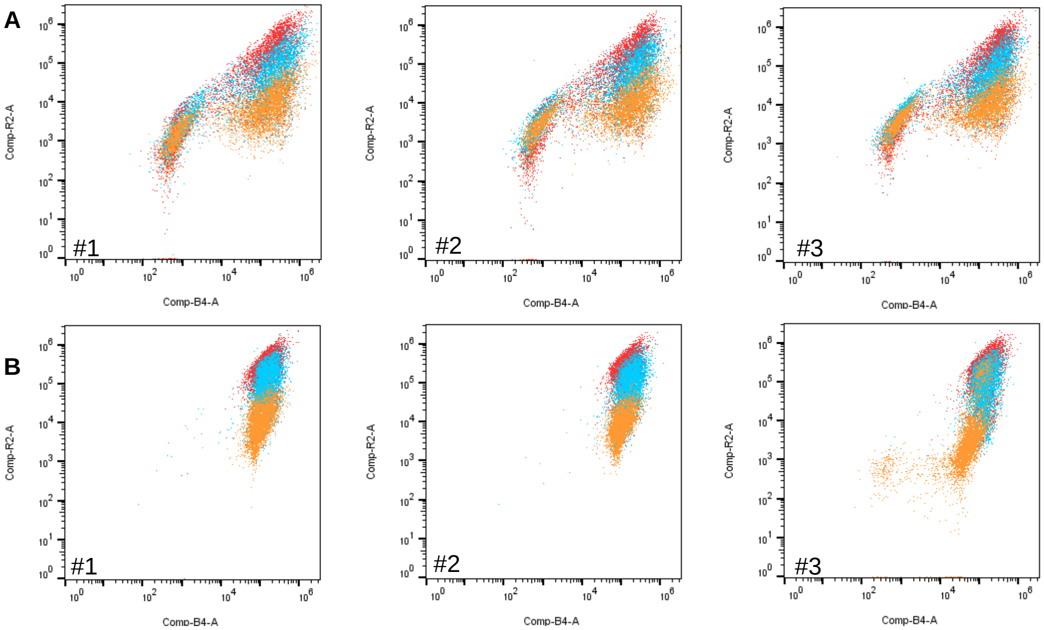


**Supplementary Figure 10. Triplicates of 3CL$^{pro}$ activity assay.** (A) 3CL$^{pro}$ in plasmid-based system. 3CL$^{pro}$ Off (red), pY2 3CL$^{pro}$ On (blue) and pY3 3CL$^{pro}$ On (orange) populations presented on anti-FLAG PE (B4-A, x-axis) v. anti-HA Alexa 647 (R2-A, y-axis) dot plots. Populations presented in log scale. (B) 3CL$^{pro}$ in integration-based system, integrated at LYS2 site. 3CL$^{pro}$ Off (red), pY2 3CL$^{pro}$ On (blue), and pY3 3CL$^{pro}$ On (orange) populations presented on anti-FLAG PE (B4-A, x-axis) v. anti-HA Alexa 647 (R2-A, y-axis) dot plots. Populations presented in log scale.

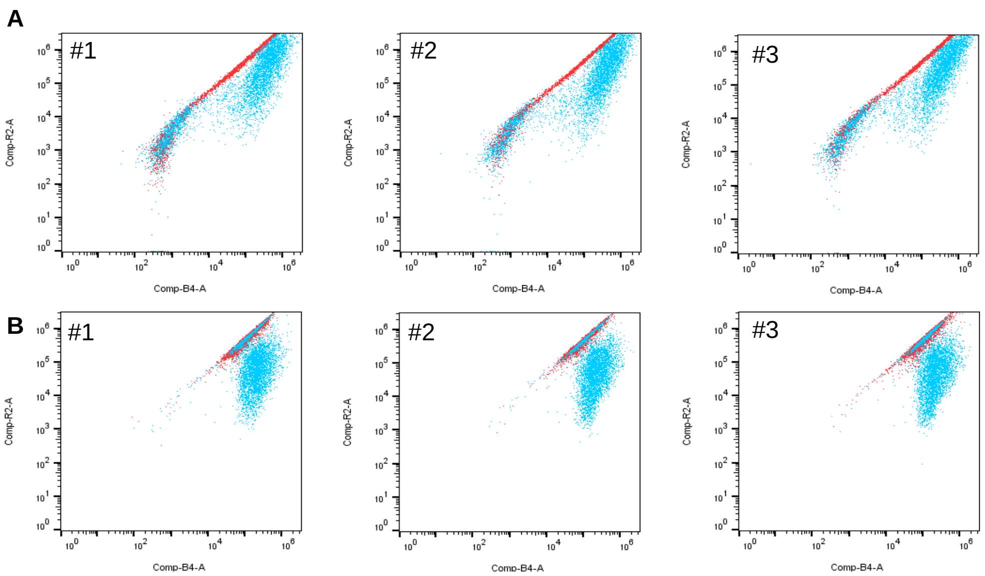

**Supplementary Figure 11. Triplicates of split-integration TEVp and TEVs systems.** (A) TEVs in plasmid-based system and TEVp integrated at the MET15 site, pY2 configuration. TEV Off (red) and TEV On (blue) populations presented on anti-FLAG PE (B4-A, x-axis) v. anti-HA Alexa 647 (R2-A, y-axis) dot plots. Populations presented in log scale. (B) TEVp in plasmid-based system and TEVs integrated at MET15 site, pY2 configuration. TEV Off (red) and TEV On (blue) populations presented on anti-FLAG PE (B4-A, x-axis) v. anti-HA Alexa 647 (R2-A, y-axis) dot plots. Populations presented in log scale.

**Supplementary Table 1. Names and sequences of oligos used in specified reactions in the manuscript.**

| Name | Oligo Sequence |
|---|---|
| TEVp-FWD | GGTCTCAAGCAAGCTTGTTTAAGGGGCCG |
| TEVp-WEH-RC | GGTCTCACTTACAGTTCATCATGTTCAAAGCTGCCATTCATGAGTTGAGTCGCTTCC |
| 3CL-FWD | GGTCTCAAGCAAGTGGTTTTAGAAAAATGGCATTCC |
| 3CL-WEH-RC | GGTCTCACTTACAGTTCATCATGTTCCCAGCTGCCTTGGAAAGTAACACCTGAGC |
| P1-FLAG-FWD | AGATACGATTATAAAGATGACGACGATAAAGGAGGTGG |
| P1-FLAG-RC | TGAGCCACCTCCTTTATCGTCGTCATCTTTATAATCGT |
| P2-TEVsub-FWD | CTCAGAAAACCTGTATTTTCAGAGCGGTGGCG |
| P2-TEVsub-RC | CTGCCGCCACCGCTCTGAAAATACAGGTTTTC |
| P2-3CLsub-FWD | CTCACTGGGCAGCGCGGTGCTGCAGAGCGGCGGTGGCG |
| P2-3CLsub-RC | CTGCCGCCACCGCCGCTCTGCAGCACCGCGCTGCCCAG |
| P3-HA-FWD | GCAGTTACCCATACGATGTTCCAGATTACGCT |
| P3-HA-RC | AACCAGCGTAATCTGGAACATCGTATGGGTAA |
| P4-FEHDEL-FWD | GGTTCGTTTGAACATGATGAACTGTAGTAA |
| P4-FEHDEL-RC | ATTGTTACTACAGTTCATCATGTTCAAACG |
| P4-WEHDEL-FWD | GGTTCGTGGGAACATGATGAACTGTAGTAA |
| P4-WEHDEL-RC | ATTGTTACTACAGTTCATCATGTTCCCACG |
| Q5-pY2-TEVsub-Removal-FWD | CACGGTTATCCACAGAATC |
| Q5-pY2-TEVsub-Removal-RC | GCTAGAATTCGTTTAAACATG |
| Q5-pY2-TEVp-Removal-FWD | GCATGTTTAAACGAATTCTAGC |
| Q5-pY2-TEVp-Removal-RC | ACTAGTGCACTGCAGTAC |
| Q5-pY2-MET- TEVsub-Removal-FWD | CAATCCATGGTCTATACATG |
| Q5-pY2-MET-TEVsub-Removal-RC | GCTAGAATTCGTTTAAACATG |
| Q5-pY2-MET-TEVp-Removal-FWD | GCATGTTTAAACGAATTCTAGC |
| Q5-pY2-MET-TEVp-Removal-FWD | ACTAGTGCACTGCAGTAC |

**Supplementary Table 2. Thermocycler protocols for specified reactions performed in the manuscript.**

| Oligo Annealing | BsaI Golden Gate | BsmBI Golden Gate | Q5 PCR | Gibson Assembly |
|---|---|---|---|---|
| 37°C 30 minutes<br>98°C 5 minutes<br>12°C 1 minute<br>Ramp<br>0.1°C/s<br>12°C Forever | 37°C 1 minute*<br>16°C 1 minute*<br>60°C 5 minutes<br>12°C Forever<br>*Cycle x30 | 42°C 1 minute<br>16°C 1 minute<br>Cycle x30<br>60°C 5 minutes<br>12°C Forever | 98°C 30 sec<br>98°C 10 sec**<br>$T_m$ 20 sec**<br>72°C 30 sec/kb**<br>72°C 2 minutes<br>4°C Forever<br>**Cycler x25 | 50°C 1 hour<br>12°C Forever |

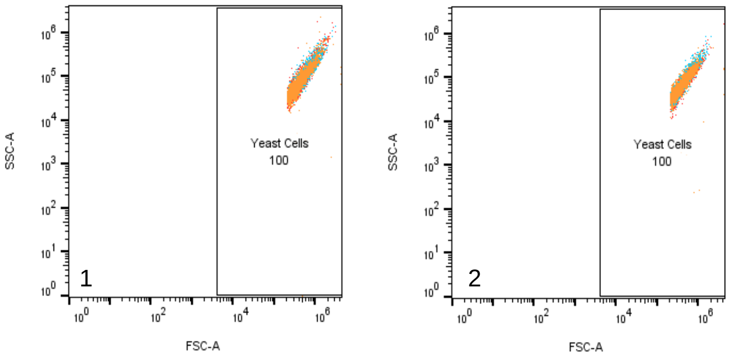


**Supplementary Figure 12. Gating strategy for capturing all yeast cells.** Representation of the rectangular gate drawn to capture all yeast cells seen on FSC-A v. SSC-A dot plots for both plasmid-based (1) and integration-based (2) systems. Sample show is #1 of the TEVp activity assay. Populations presented in log scale.

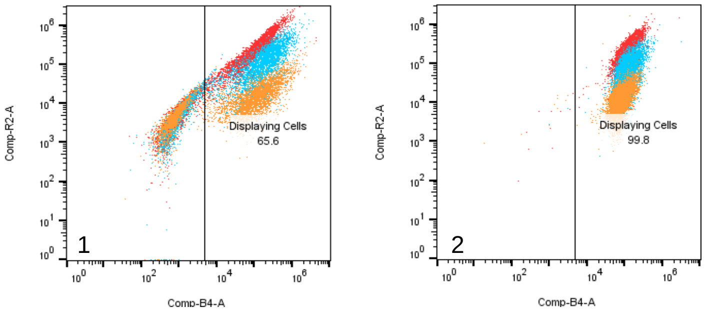


**Supplementary Figure 13. Gating strategy for capturing displaying cells.** Representation of the rectangular gate drawn to capture displaying cells seen on anti-FLAG PE (B4-A) v. anti-HA Alexa 647 (B2-A) dot plots for both plasmid-based (1) and integration-based (2) systems. Sample show is #1 of the TEVp activity assay. Populations presented in log scale.

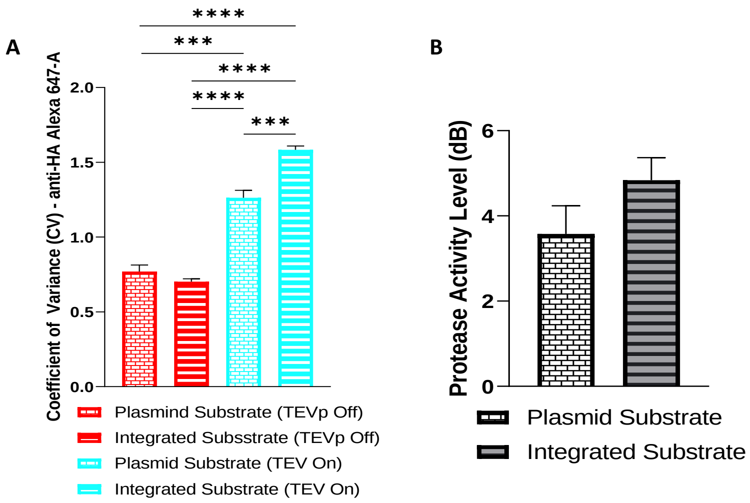


**Supplementary Figure 14. Population spread and signal-to-noise ratios of split-integration TEVp and TEVs systems.** (A) Coefficient of Variance ($CV_{HA}$) of the of the two split-cassette configurations was calculated on the displaying yeast cells and the variability of anti-HA Alexa 647-A fluorescence signal across the cell populations. $CV_{HA}$, a quantification of population spread, is dependent on the uniform expression of the genes that translate to a signal being observed. (B) Protease activity level ($D_{dB,HA}$) calculated using Equation 1 and represented in units of dB. No significance was observed between activity levels observed in both split-cassette systems. Statistical significance between populations was determined by multiple unpaired t-tests. *p ≤0.05, **p ≤0.01, ***p ≤0.001, ****p ≤0.0001.
